# Algal symbioses with fire corals demonstrate host genotype specificity and niche adaptation at subspecies resolution

**DOI:** 10.1101/2023.04.03.535406

**Authors:** Caroline E Dubé, Benjamin CC Hume, Emilie Boissin, Alexandre Mercière, Chloé A-F Bourmaud, Maren Ziegler, Christian R Voolstra

## Abstract

Corals share an intimate relationship with photosynthetic dinoflagellates that contribute to the biology of the emerging metaorganism. While many coral-algal associations exhibit high host fidelity, the extent of this specificity under environmental change remains to be fully understood and is a prerequisite to forecasting the adaptive potential of this obligate symbiosis. Here, we disentangled the contribution of host genotype and environment on governing coral-algae associations by working at subspecies resolution. We used fine-scale genotyping of algal symbionts from 198 fire coral colonies (*Millepora* cf. *platyphylla)* that map to ten distinct sexually produced clonal host genotypes across three environmentally distinct reef habitats. Based on microalgal ITS2 genotyping, we show that algal-host specificity extends down to the Symbiodiniaceae subspecies level in a natural reef environment. Closely related *Symbiodinium* (A7)*-*dominated algal assemblages almost perfectly mapped to fire coral host genotype. Furthermore, identification of host genotype- and habitat-specific *Symbiodinium* alga suggest the presence of algal phenotypic diversity even at this taxonomic resolution (i.e., within *Symbiodinium* A7), which may aid environmental niche adaptation of the metaorganism. Our results suggest that the here-identified *Millepora*-*Symbiodinium* associations are co-evolved to match their prevailing environment. Thus, despite the presence of rarer host generalist *Cladocopium* algae, scope for environmentally induced modification of the cnidarian-algal association is likely constrained by host genotype.

## Introduction

Symbiotic microorganisms are increasingly recognized as foundational components of animal hosts, thereby impacting the ecology and physiology of their hosts^1^. Reef-building corals (among other cnidarians) form an obligate endosymbiosis with dinoflagellate algae of the family Symbiodiniaceae^2^. The algal symbionts satisfy up to 95% of their host energetic requirements^3, 4^, while the coral host provides a protected light-rich environment and supply of nutrients^5^. This symbiosis creates the foundation of coral reef ecosystems and is one of the oldest, most widespread, and ecologically successful known mutualisms found in nature^6^. The rapid pace of environmental change caused by climate warming and other stressors is now posing a serious threat to coral reef ecosystems and has been linked to numerous occurrences of coral bleaching (severe loss of algal symbionts under stressful conditions)^7, 8^.

Algal identity is an important factor contributing to variability of stress tolerance in corals^9–12^. This variability is likely in part due to the considerable genetic and phenotypic diversity found within each of the most common coral-associating algal genera (*Symbiodinium*, *Breviolum*, *Cladocopium*, and *Durusdinium*^13^) characterized by their own physiological and ecological traits^2, 14–16^. Such phenotypic diversity means that certain assemblages are better suited for specific prevailing environments than others, as evidenced by the observed environmental structuring of coral-algal associations^17–20^. Changing algal associations may therefore act as a mechanism of ecological adaptation/acclimatization. However, the degree to which corals are able to modify their dominant algal symbiont association appears dependent on the host and algal genotype. Notably, host and symbiont specialists and generalists exist^18, 21–23^. There are many examples of coral colonies exhibiting fixed algal associations under environmental change ^24–27^. Conversely, environmentally induced changes from a predominant alga of one genus to a different genus are documented and are exemplified by the appearance of the stress-tolerant *Durusdinum trenchii*^12, 28^. However, those changes are typically transient and revert back to the original association when stress subsides^14, 27, 29^. These contrasting examples highlight that the identity of a given coral-algal assemblage is therefore a consequence of host genotype and environment.

Disentangling the influence of host genotype and environment on coral-algal associations is challenging, since host genotype needs to be controlled for in the sampling design and/or experimental setup. Further, methods used for algal genotyping must be able to resolve differences at the desired taxonomic resolution^30^. To date, host genotype has often been controlled for at the species level or even higher (e.g., genus), demonstrating host-specificity in most cases^31–33^. However, the relevance of any conclusions on host-specificity by such studies is limited to the resolution at which the host has been resolved. Thus, to investigate whether host fidelity can also be observed below the species level (i.e., genotype), subspecies resolution of the host is required. Further, individuals harboring the same genotype must be distributed and assessed across different environments to disentangle the relative contribution of host genotype and environment to prevalent host-algal associations. This has been addressed by investigating algal associations of clonal coral replicates transplanted across environments or captively reared, showing that Symbiodiniaceae assemblages are highly host genotype-specific^21, 34, 35^.

Here, we ITS2-genotyped the algal assemblage of 198 hydrocoral samples (*Millepora* cf. *platyphylla*) previously characterized to belong to ten different clonal genotypes with members of each genotype distributed across at least two environmentally distinct reef habitats. In doing so, we could assess algal associations across environments in dependence on host genotype in a natural population at a subspecies level. We demonstrate that algal-host specificity extends down to the subspecies level of host and symbiont. This argues for phenotypic diversity even between highly similar algae and implies that host-algal symbiotic associations are likely constrained at the level of individual genotypes.

## Results

### Fire coral clones are exposed to distinct environmental conditions

To discriminate the relative contribution of host genotype and environment on structuring coral-algal associations, we used microalgal ITS2 genotyping from ten clonal genotypes of the fire coral *M.* cf. *platyphylla* distributed across three reef habitats (mid slope, upper slope, back reef) that exhibit contrasting environmental conditions, as shown by temperature and light data obtained from *in situ* deployed loggers (Fig. 1). Although temperature profiles showed a similar daily mean water temperature across the three habitats (back reef, BR: 26.89 ± 0.07°C, upper slope, UP: 26.79 ± 0.05°C, mid slope, MD: 26.74 ± 0.04°C), they exhibited a three- to four-fold greater diel amplitude at the back reef (1.37 ± 0.43°C) compared to both fore reef habitats (UP: 0.45 ± 0.18°C, MD: 0.34 ± 0.15°C). Daily maximum temperatures were significantly higher at the back reef (BR: 27.71 ± 0.08°C, UP: 27.05 ± 0.06°C, MD: 26.92 ± 0.05°C, Kruskal-Wallis, P < 0.001) (Supplementary Fig. 2). Light intensity profiles revealed a similar pattern with significantly higher daily mean and maximum irradiance at the back reef (446.28 ± 20.99 and 2 271.69 ± 63.80 µmol photons/m^2^/s, respectively) compared to the upper slope (266.57 ± 13.36 and 1 371.83 ± 46.54 µmol photons/m^2^/s) and the mid slope (137.50 ± 6.82 and 726.38 ± 22.55 µmol photons/m^2^/s, Kruskal-Wallis, P < 0.001) (Supplementary Fig. 2).

**Figure 1.**
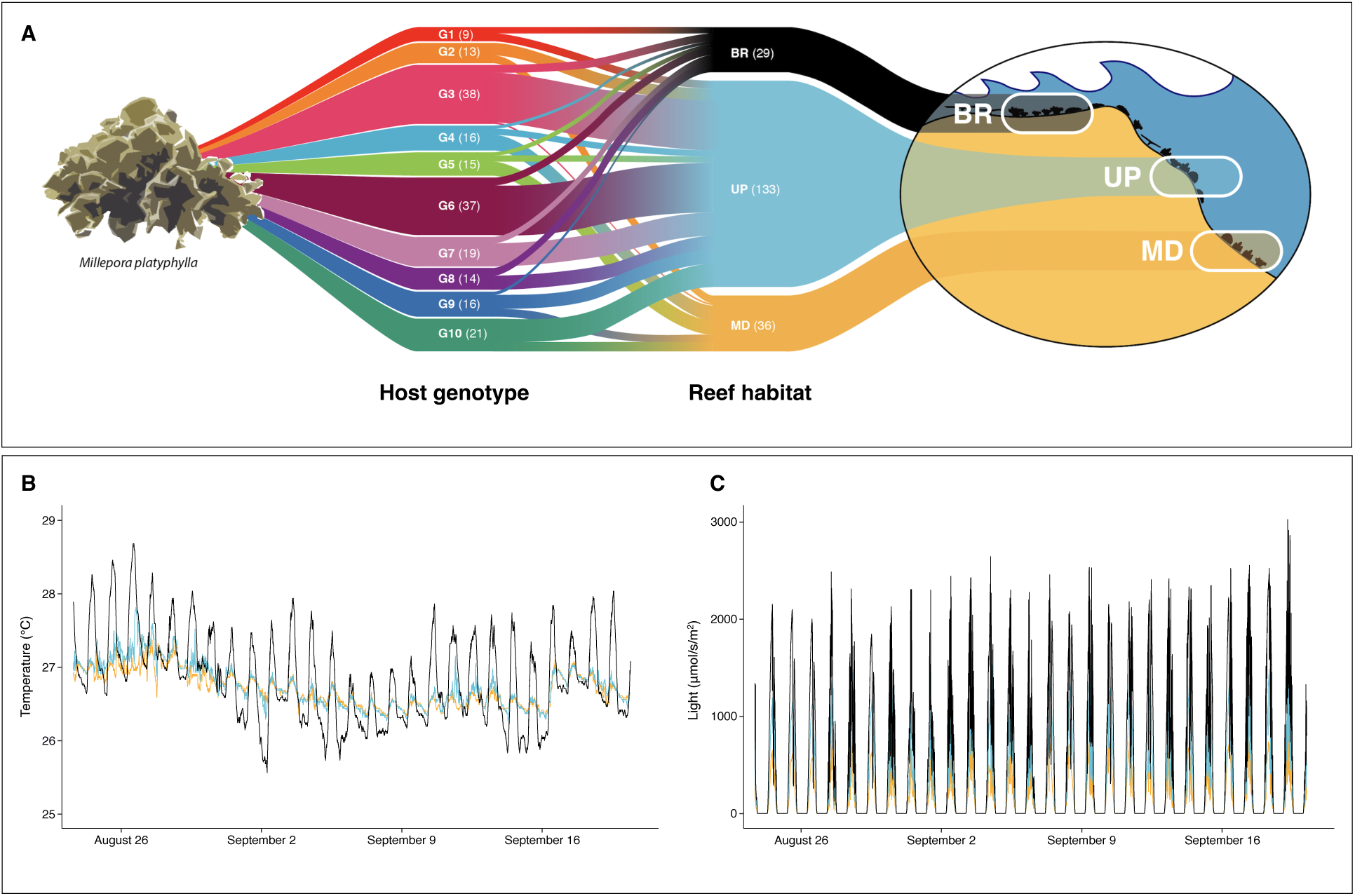
Schematic overview of the sampling design. **A** On the left of the alluvial plot, the ten sampled clonal genotypes are shown (G1 to G10) with clones naturally distributed across the reef habitats on the north shore of Moorea, French Polynesia: the back reef (BR at < 1m depth), the upper slope (UP at 6 m), and the mid slope (MD at 12 m), which are illustrated on the right. Each genotype and habitat are represented by a unique color with faded colors towards the middle where clones of a given genotype are aligned with the habitat in which they were found. The thickness of each branch corresponds to the number of replicates per genotype and habitat shown in parentheses. In the middle, the thickness of each branch corresponds to the abundance of clones (n) for each genotype in each of the three habitats. **B** Temperature profiles from each of the three habitats (BR = black, UP = blue, MD = gold), based on 1 440 data points collected daily for a period of 29 days in the months of August to September. **C** Light profiles at each of the three habitats based on 960 data points collected daily for a period of 29 days in the months of August to September.

### Symbiodiniaceae ITS2 sequence assemblage

From the 198 fire coral samples, a total of 419 post-MED Symbiodiniaceae sequences were retrieved from the sequencing of the ITS2 amplicons that belonged to six genera: *Symbiodinium* (n = 107), *Cladocopium* (n = 276), *Durusdinium* (n = 2), *Gerakladium* (n = 27), *Fugacium* (n = 4), and *Halluxium* (n = 3) (corresponding to previously described clades A, C, D, G, F, and H, respectively)^4^ (Supplementary Data 2). Eight fire coral samples contained sequences representing at least three genera, 47 contained sequences representing two genera (44 associations of *Symbiodinium*-*Cladocopium* and 3 of *Symbiodinium*-*Gerakladium*), and 143 samples contained sequences from a single genus (of which all were from *Symbiodinium*). The majority of sequences were from *Symbiodinium* (Fig. 2A), accounting for 99.8% of the total Symbiodiniaceae reads and for at least 92% of reads per sample. Six (out of 107) *Symbiodinium* sequences occurred in all 198 samples: A7 (the most abundant across all samples ranging from 30 to 59% of reads per sample), A16d (7–27%), A3 (1–9%), A3a (1–8%), A7b (1–6%), and A7a (1–5%) (Fig. 2B). *Cladocopium* sequences were present in 52 of the 198 samples (26%) and in those samples accounted for at most 8% of reads. The two most common *Cladocopium* sequences, C15 and C116, occurred at a maximum relative abundance of 5% and 3% of Symbiodiniaceae reads, respectively. Differences in the rDNA copy number between different taxa may confound the translation of Symbiodiniaceae ITS2 counts to actual relative abundances of algal cells – especially for inter-genera comparisons. Current estimates suggest *Cladocopium* has a higher copy number than *Symbiodinium* (means of 2119 ± 217 and 1721 ± 125, taken respectively from Saad et al.^36^). As such, the greater abundance of *Symbiodinium* compared to *Cladocopium* ITS2 reads observed in our samples likely relate to a higher abundance of *Symbiodinium* cells.

**Figure 2.**
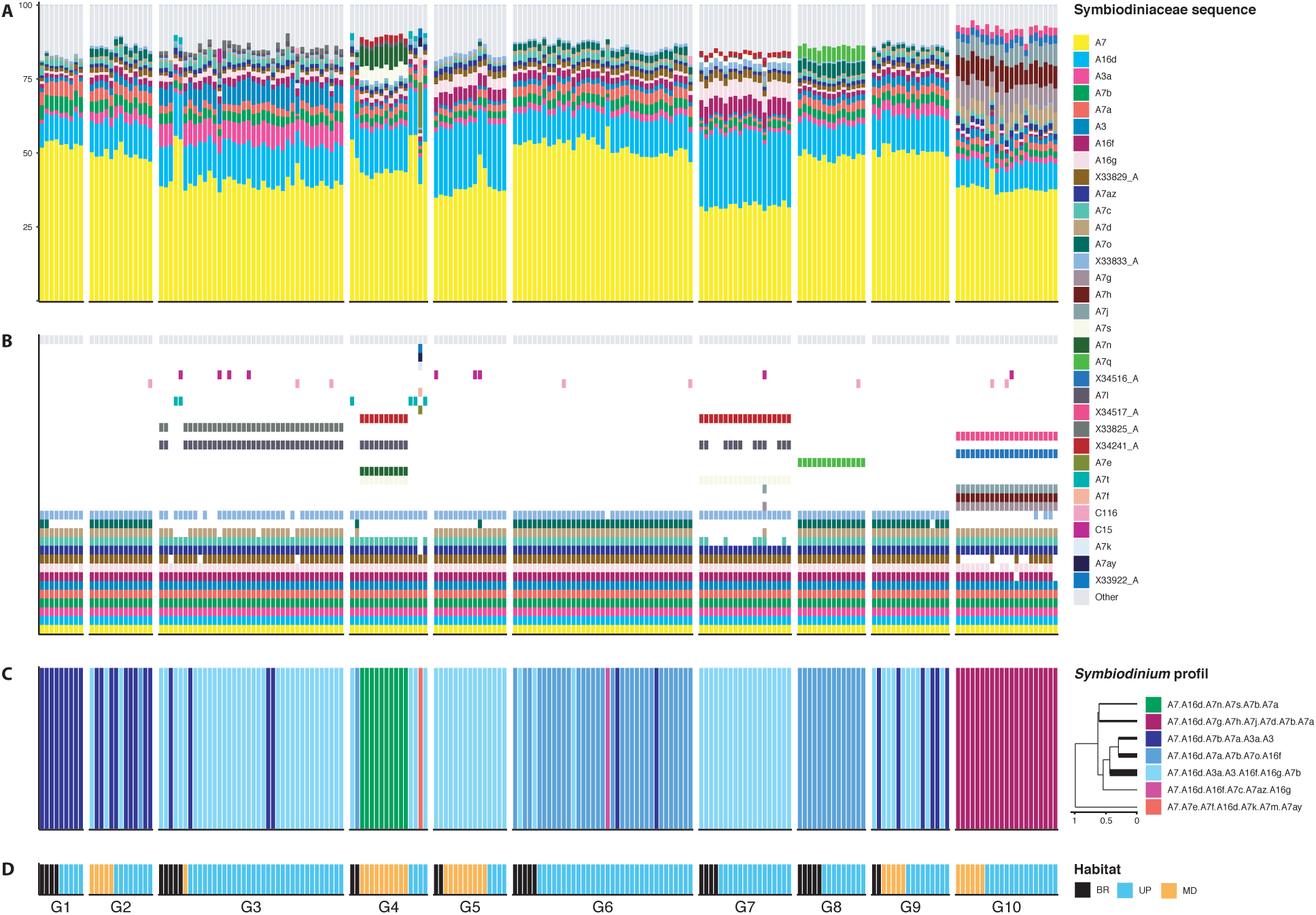
Symbiodiniaceae community composition of 10 fire coral clones across 3 distinct reef habitats. In each of the main figure sections (A-D), every sample is represented as a single column with samples grouped by host genotype and reef habitat denoted in **D. A** Relative abundance of post-MED ITS2 sequences (as a proxy for Symbiodiniaceae taxonomic composition). **B** Presence or absence of ITS2 sequences regardless of abundance. Missing bars represent sequences not present in the sample. For both **A** and **B**, sequences are ordered from bottom to top by overall relative abundance across all study samples. Sequences that were found in all samples with a relative abundance > 2.5% are annotated in the legend. All other sequences are grouped as ‘Other’ and are shown in light grey. **C** Predicted ITS2 type profiles (as a proxy for Symbiodiniaceae genotypes). Profiles are annotated in the dendrogram on the bottom right. The dendrogram shows the hierarchical clustering of Bray-Curtis-derived between-profile distances calculated from the average relative abundance of defining intragenomic variants for each of the profiles (from 0 to 1). Thickness of leaves corresponds to the abundance of the respective *Symbiodinium* type profile. **D** Host genotype and reef habitat of origin. Clonal genotypes are numbered from G1 to G10 and each genotype contains multiple samples (n) that occur naturally in at least two of the three surveyed habitats (BR = black, UP = blue, MD = gold).

### Symbiodiniaceae ITS2 profiles

SymPortal identified a total of 18 ITS2 profiles (used as a proxy for distinct Symbiodiniaceae genotypes, i.e. genetically distinct entities putatively belonging to one or more species within each of the genera) defined by 56 DIVs that belonged to three Symbiodiniaceae genera: *Symbiodinium* (7 profiles), *Cladocopium* (10 profiles), and \*Gerakladium* (1 profile; Supplementary Data 3). The one *Gerakladium* profile was not considered further in the analysis as it only occurred in a single sample at an abundance < 0.01%. The fire corals in this study were dominated by *Symbiodinium* profiles with each of the 198 samples predicted to contain only one of the identified *Symbiodinium* profiles (Fig. 2C). *Symbiodinium* profiles were all highly similar with each of the above mentioned six common ITS2 sequences (A7, A16d, A3, A3a, A7b, and A7a) found in at least two of the profiles and with ITS2 sequences A7 and A16d found in all profiles. The profiles were unevenly distributed across samples with three of the most similar profiles found in 82% of the samples (Fig. 2C). In order of abundance, the five most abundant ITS2 profiles were found in 88, 47, 30, 21, and 10 of the fire coral samples with the remaining two ITS2 profiles (i.e., putative genotypes) found in only one fire coral sample each (Fig. 2C). Of the five most abundant *Symbiodinium* profiles found in this study (Supplementary Data 3), only one was found in other samples in the SymPortal remote database. These were two *Millepora* cf. *platyphylla* samples collected in the same reef system as this study, but three years later (November 2016), as part of the *Tara* Pacific Expedition^37^. Finally, the two ITS2 profiles found in only one sample from this study were found in 43 and 2 samples from the SymPortal database also collected as part of the *Tara* Pacific Expedition in a separate sampling effort by E. Boissin in November 2015 (*M.* cf. *platyphylla*; unpublished). While the majority of *Tara* Pacific samples were collected in the same reef as this study’s samples, 6 of the 43 were collected from reefs off the coast of Fiji over 3000 km away.

Ten unique *Cladocopium* profiles were found in 15 of the samples and were always below 9% relative abundance (Supplementary Fig. 3). Of these ten ITS2 profiles, six were defined by only 1 or 2 DIVs indicating that SymPortal likely did not sample the *Cladocopium* genotypes they represent a sufficient number of times (i.e., they were not present in a sufficient number of samples) to be able to generate profiles with a greater number of DIVs that would be more specific representatives of these genotypes. However, four of the profiles (characterized by the C91, C15, C1, and C116 sequences as the respective most abundant sequences) were defined by at least 4 DIVs suggesting that they were likely specific representatives of the underlying *Cladocopium* genotypes. These *Cladocopium* profiles were found in 4, 3, 1, and 1 of the samples from this study (Supplementary Data 3) and in 0, 2, 96, and 1 samples from other studies in the remote SymPortal database. Unfortunately, only two of these samples from the other studies (both containing the C15 profile) are publicly available. These two samples derive from *Stylophora pistillata*, one from the northern Red Sea and one from the central Red Sea^38^.

### High fidelity of *Millepora* host genotypes and algal assemblages

The explanatory power of host genotype and reef habitat variables on the variation in Symbiodiniaceae community composition was assessed. Differences in ITS2 sequence assemblage (i.e., total taxonomic composition) were predominantly explained by host genotype (two-way PERMANOVA, *F* = 119.5, *R*^2^ = 0.832, *P* < 0.001), but also by the interaction of host genotype and reef habitat (*F* = 3.6, *R*^2^ = 0.034, *P* < 0.001). Accordingly, samples demonstrated a discrete grouping by host genotype in the PCoA with only three exceptions: G4 samples grouped by host genotype and habitat, two G3 samples grouped with G4 back reef and upper slope samples and with one G4 upper slope sample grouping reciprocally, and the G6 and G8 samples overlapped in their ordination coordinates in the first three PCs (Fig. 3). However, G6 and G8 samples were clearly differentiated by the presence of the A7q ITS2 sequence in all of only the G8 samples (Fig. 2B). Genotype-specific *Symbiodinium* sequences were also identified (29 in total and ranging from 1 to 8 per genotype; Supplementary Data 2). Of note, reef habitat was also significant in explaining ITS2 sequence assemblage (i.e., total taxonomic composition) although with a much weaker explanatory power than host genotype (two-way PERMANOVA result; *F* = 5.574, *R*^2^ = 0.009, *P* < 0.001).

**Figure 3.**
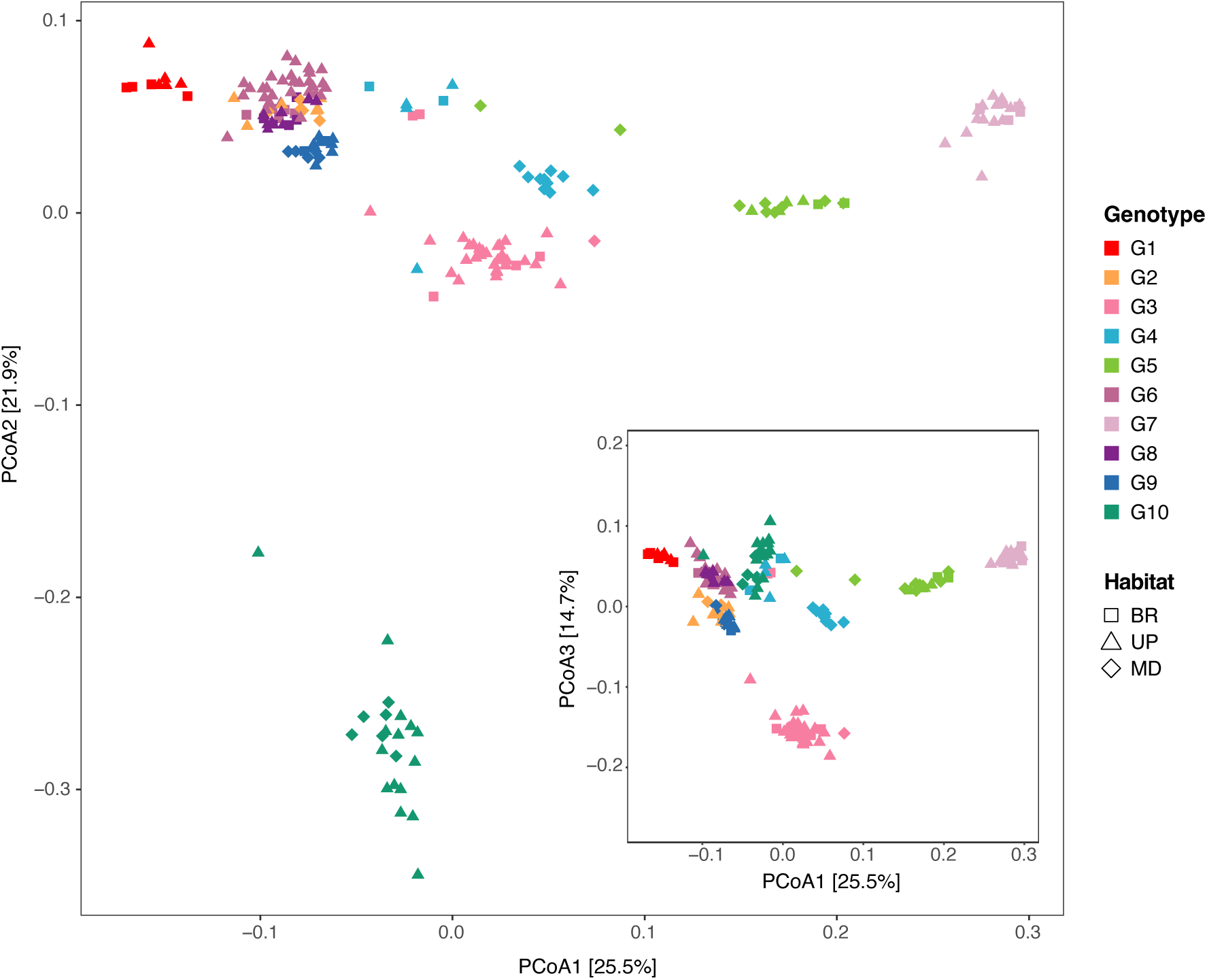
Contribution of host genotype and habitat to *Symbiodinium* assemblage. PCoA of square root-transformed relative abundances of *Symbiodinium* (A-)sequences constructed using Bray-Curtis distances. Each dot represents a colony, colors denote the genotype, and symbols denote the reef habitat.

In support of the ITS2 sequence-based analysis, the ITS2 profile analysis predicted discrete profiles for all G10 samples (genotype-specific) and the G4 mid slope samples (genotype- and habitat-specific with the unique DIV A7n) (Fig. 2B and 2C). However, other abundant algal profiles were present in multiple host genotypes. The clear (host genotype-specific ITS2 sequences) and consistent (found across multiple samples from each host genotype) pattern of algal assemblage differentiation between members of different host genotypes seen in the sequence-based analysis suggests that the process of ITS2 profile prediction (in SymPortal) did not resolve the likely host-specific algal genotypes.

Mantel tests further confirmed host genotype specific algal assemblages by revealing a significant positive correlation between Symbiodiniaceae community composition dissimilarities and host genetic distances (*r* = 0.3258, *P* < 0.001). In other words, genetically more distant host genotypes were associated with more dissimilar Symbiodiniaceae communities (Supplementary Fig. 4). For instance, genotypes G5 and G7 (34 samples) were closely related genetically (Supplementary Fig. 1) and characterized by similar Symbiodiniaceae communities (Fig. 3), including association with the same *Symbiodinium* profiles (Fig. 2C). Correspondingly, genetically divergent genotype G10 (Supplementary Fig. 1) was uniquely and solely associated with one ITS2 profile that was genetically distinct based on average relative abundance of defining intragenomic variants (Fig. 2C).

### Symbiodiniaceae community dissimilarity correlates with fine-scale spatial distance of *Millepora* colonies

The sampling design with georeferenced clonal colonies enabled us to investigate Symbiodiniaceae community composition of fire coral genotypes across small spatial scales. From ten mantel tests performed for each of the ten host genotypes, a significant positive correlation between the Symbiodiniaceae community composition dissimilarity and the spatial distance between clones across reef habitats (Euclidean distance in meters based on x and y spatial coordinates of each clone) was demonstrated for five of the genotypes (G1, G3, G4, G6, G8). In other words, more spatially distant clones were associated with more dissimilar Symbiodiniaceae communities (Supplementary Table 1). However, because these tests contained clones distributed in multiple habitats, the effect of spatial distance is confounded with the effects of reef habitat. To separate the effect of spatial distance from reef habitat, we performed the same correlations, but for distinct genotype-habitat combinations (Supplementary Table 1). Significant positive correlations were returned for three of the twenty tests (Fig. 4) demonstrating fine-scale spatial variation (distances ranging from 0.07 to 20 meters) in algal assemblage even between clones of the same habitat.

**Figure 4.**
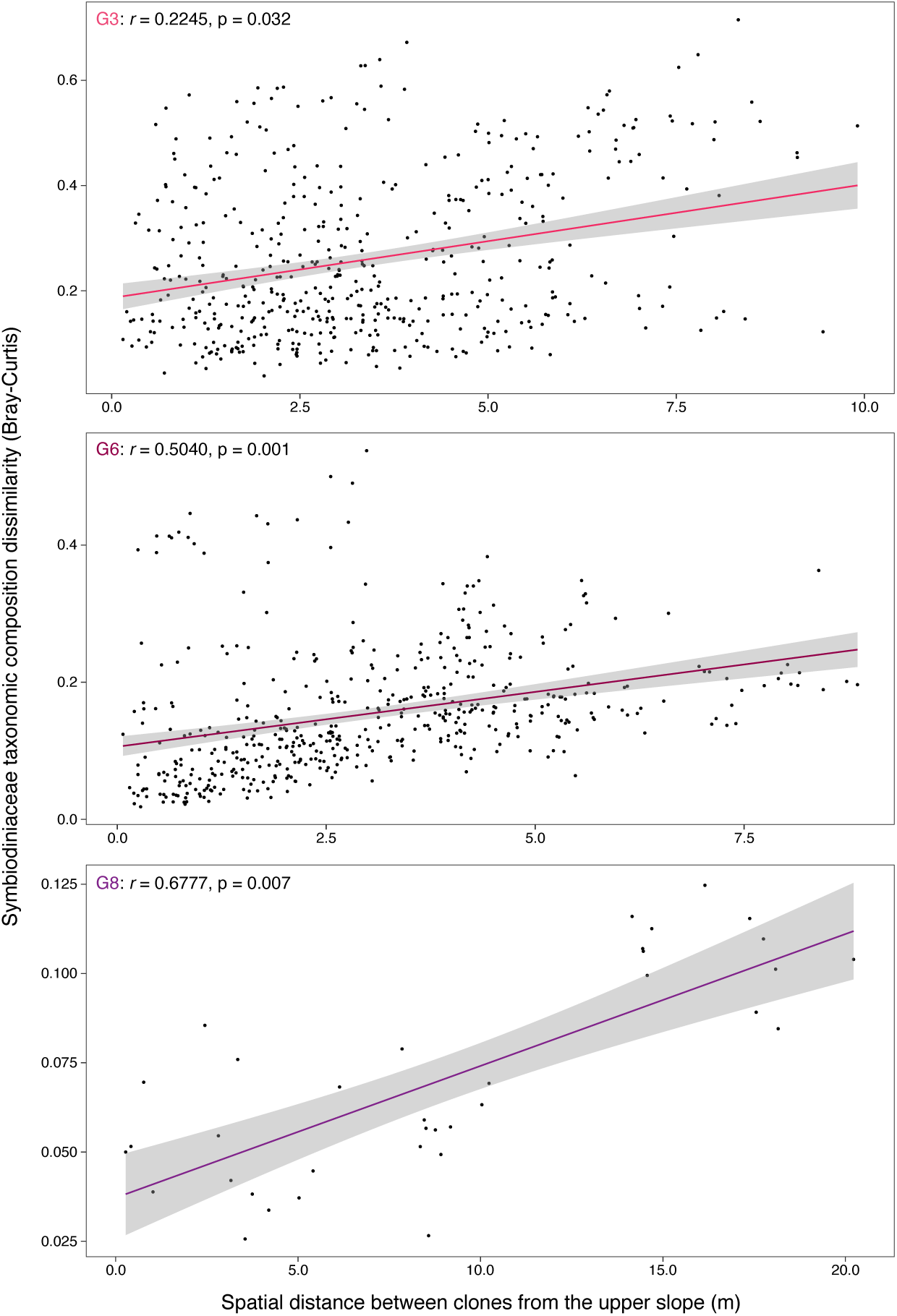
Relationship between *Symbiodinium* ITS2 sequence assemblage dissimilarity and spatial distance of clones in reef habitats. Positive correlations between relative abundances of *Symbiodinium* (A-)sequences constructed using Bray-Curtis and host spatial distances (Euclidean matrix in meters). Only clones of the genotypes G3, G6, and G8 originating from the upper slope are shown, with their respective Spearman’s Rank correlation coefficient alongside p-values estimated with a Mantel test. Correlations for the other genotype x habitat combinations were not significant (Supplementary Table 1).

## Discussion

Identifying coral-algal symbiosis specificity and the extent to which the host and its symbionts co-diversify in response to environmental change is essential for forecasting how this obligate symbiosis may promote or restrict the potential for coral adaptation under climate change^39^. Numerous studies have convincingly demonstrated the importance of host identity on Symbiodiniaceae assemblage^31–33^, including environmentally induced variations in symbiont composition^40–42^. Yet studies that are able to disentangle the relative influence of animal host and environment on partner pairing dynamics below the species level by controlling for host genotype are missing at large. Here, we assessed the extent of environmental-induced variations in Symbiodiniaceae community composition in reef-building corals, while controlling for host genotype at a clonal level without the manipulation of corals through transplantation or captive rearing.

Species definitions within the Symbiodiniaceae are becoming more common, and recent definitions are starting to formally recognize the large extant algal richness. However, the majority of coral-associating algal symbionts still lack a formal definition^30, 43^. Unfortunately, many of the existing formal taxonomic descriptions lack information regarding ITS2 sequences. For those definitions that do exist, they are often characterized by on average one to three dominant ITS2 sequences (e.g., one or two dominant sequences for *Symbiodinium*^44, 45^ and *Cladocopium*^46, 47^, and one to three for *Durusdinium*^48^). Thus, characterization of our samples goes far beyond these two-sequence definitions.

Our study revealed that algal assemblage maps almost perfectly to host genotype, at a subspecies host and algal resolution (Fig. 2 and 3). While we observed disagreement between the ITS2 sequence- and profile-based approaches, this is likely due to the lack of resolving power at this fine-scale level by SymPortal. Rather, our results suggest that a combination of sequence-based and profile-based analyses as employed here in some cases are preferable to yield the resolutive power needed to identify patterns at this fine-scale resolution. Here we can show that the importance of host identity in structuring coral-algal assemblages extends to the level of subspecies genotype. The strong host genotype fidelity observed is consistent with previous reports showing that host identity largely determines the physiological response of the symbiont, as well as the stability and functionality of the symbiosis to changing environmental conditions^35, 49^. Notably, variations in the physiological response to increased temperature within a single Symbiodiniaceae species or clonal host-symbiont combinations have been previously described in temperate corals^16^ and Aiptasia^49^, but are rare for tropical corals^50^. Our study also showed that host genetic similarity aligns with Symbiodiniaceae assemblage similarity, irrespective of host habitat, which is often expected for species that acquire their symbionts via vertical transmission^51^. Unlike many broadcast spawning corals, *Millepora* hydrocorals can vertically transmit their algal symbionts to their eggs^52^, thus exhibiting a mixed mode of symbiont acquisition. This may explain, at least to some degree, the strong host genotype specificity observed for *Millepora*-associated *Symbiodinium* algae. The establishment and maintenance of such specific coral-algae associations likely involve cell-signaling and other inter-partner recognition mechanisms, so that intracellular symbionts may co-evolve with the host to become particularly well suited to meet the metabolic demands of their host’s micro-environment. *Symbiodinium* characterized with their most abundant ITS2 sequences of A7 (hereafter simply referred to as *Symbiodinium* A7, and a format we will use for describing other alga associated to specific ITS2 sequences) is known to associate with hydrozoans, but not with anthozoans in the Pacific Ocean^53^. This suggests that this alga might be highly host-specialized due to co-evolutionary processes with *Millepora* hydrocorals. Many members of the *Symbiodinium* genus are hosted by corals found in variable environments^54^ and are often associated with increased tolerance to high irradiance and temperature^38, 45^. Accordingly, fire corals have been described as thermotolerant species during climate-driven marine heat waves in French Polynesia, although many colonies bleached and died from the recent massive bleaching event of 2019^52^. This thermal tolerance might be linked to their partnership with specific *Symbiodinium* algae, as previously reported for bleaching-resistant Caribbean *Millepora* species^55^.

Despite a symbiont community dominated by *Symbiodinium*, our results revealed that *M.* cf. *platyphylla* can also associate with a variety of *Cladocopium* algae, as seen previously in the Pacific Ocean^31, 56^. Samples from our study associated with *Cladocopium* C91, C15, C1, and C116, while *Millepora* in the wider Pacific were reported to associate with C57 and C66. The presence of these rarer *Cladocopium* algae may be explained by *Millepora*’s uncommon life history trait combination of broadcast spawning of medusoids (which then release sexual gametes) with a (non-strictly) vertically transmitting symbiosis. The increased duration of time larvae spend in the water column may allow for an increased exposure to exogenous algal symbionts compared to strictly vertically transmitting symbiosis (typical of brooders). If host uptake mechanisms allow, this may enable the larvae to associate with symbionts from the water column that are not present in the maternal colony^57^. Of particular interest is the apparent cosmopolitan and host-generalist nature of at least one (a *Cladocopium* C15), but possibly more of the *Cladocopium* algae that *Millepora* associated with in this study (the genotypes represented by the *Cladocopium* C1 and C116 profiles were found in additional samples from the SymPortal database). These associations represent a contrast to the apparent host-specific *Symbiodinium* algae that otherwise dominated the *Millepora* assemblages. Association with *Cladocopium* algae may allow fire corals to respond more promptly to changes in environmental conditions across reefs through shuffling of background species^19, 58, 59^ and an increase in the number of different genera hosted by coral has been posited to be a possible response to climatic perturbations^38^.

Although host genotype was the main factor explaining variance in Symbiodiniaceae assemblage structuring, the interaction of reef habitat with host genotype and reef habitat alone returned significant results in statistical testing, although with a much-reduced strength of correlation. In addition, our fine-scale mapping of fire corals on the reef substratum revealed that for three of the genotypes (Fig. 4), colonies separated by smaller spatial distances associate with more similar Symbiodiniaceae communities compared to more distant colonies. This suggests that host micro-habitat may influence coral-algal structuring, adding to the potential for fine-scale environmentally induced variations (i.e., isolation by distance (IBD) within reef habitat). In line with this environmental effect, we identified one habitat-specific (i.e., endemic) *Symbiodinium* profile (with the unique DIV A7n) in all clones belonging to the host genotype G4 from the mid slope, while those on the upper slope and back reef were associated with other *Symbiodinium* profiles. This result suggests a phenotypic diversity within these closely related algae upon which natural selection may act, similar to previously reported within species phenotypic diversity but at a more fine-scale resolution^56, 60^, enabling niche adaptation of the coral-algal association. Specifically, this habitat-endemic *Symbiodinium* genotype may be functionally distinct and adapted to the environmental conditions characterizing the mid slope habitat, e.g. a lower irradiance, though detailed characterization of this functional diversity to understand its potential role as a substrate for ecological adaptation requires dedicated investigations characterizing the physiological performance of these host-algal associations, such as light harvesting, carbon production, and nutrient translocation. Interestingly, no *Symbiodinium* genotypes were uniquely associated with the back reef, even though corals are exposed to warmer temperature and high irradiance than their reef slope counterparts. Light and temperature are known to exert a strong selection pressure on dinoflagellate algae, often leading to the zonation of Symbiodiniaceae populations between reef habitats^42, 61, 62^. This was not the case in our study, suggesting that such zonation may depend on the host species or geographic region.

## Conclusion

With reef ecosystems under stress from climate change on a global scale, it is important to consider how the host and environmental structuring we have identified here will impact the adaptive potential of *Millepora* corals to future environmental change^39^. Our data demonstrates high host-fidelity in the *Millepora*-*Symbiodinium* associations and a limited effect of the environment. The relevance of these findings may be viewed from two contrasting perspectives. It may be that the individual algal genotypes that are found across the different reef habitats are sufficiently phenotypically plastic to form successful associations with their genotype-specific coral hosts (i.e., able to survive in the long-term). Conversely, the high fidelity of the *Millepora*-*Symbiodinium* associations may be viewed as limiting, with the coral host unable to maintain alternative associations over the long term, either due to a lack of alternative host-specific symbionts, or due to host-generalist symbionts being incompatible. The former scenario is more parsimonious given the documented selectable phenotypic diversity within the *Millepora*-associating *Symbiodinium* algae and a weak but present host-independent effect of the environment on algal assemblage (indicating the algal assemblage may be modified according to environmental forcings at least to some degree). Furthermore, our identification of (possibly multiple) cosmopolitan, host generalist *Cladocopium* algae suggest the ability of the host to associate with generalist symbionts. However, their low overall abundance indicates either that these symbionts are outperformed by host-specific algal associations or that these symbionts are poorly suited to the prevailing environment. Taken together, our results suggest that although the current *Millepora-Symbiodinium* associations may well suit their environment, there is scope for putative adaptive, environmentally induced algal assemblage modification, selecting upon the extant host-specific co-evolved *Symbiodinium* diversity.

## Methods

### Fire coral sampling

Between May and September 2013, 198 fragments of the fire coral *M.* cf. *platyphylla* were collected on the north shore of Moorea Island, French Polynesia (17.5267 S, 149.8348 W), from three reef habitats, located in front of Papetoai village, that are in close proximity to each other, the mid slope (13 m depth), upper slope (6 m depth), and back reef (< 1 m depth), but presumably exhibit contrasting environmental conditions, including varying exposure to temperature, solar irradiance, partial pressure of carbon dioxide, and disturbances^63–65^. Our sampling design is described in detail in Dubé et al.^66^, where fire coral colonies were sampled to investigate the clonal structure between habitats on a barrier reef system. Briefly, three 300 m-long by 10 m-wide belt transects were laid over the reef within each habitat, parallel to the shore. All colonies of *M.* cf. *platyphylla* were georeferenced by determining their position along the transect-line (0 to 300 m) and straight-line distance from each side of the transect (0 to 5 m, totalling 10 m). From these measures, each colony was mapped with x and y coordinates. The colony size (projected surface) of each colony was estimated (in cm^2^) from 2D photographs using ImageJ 1.4f^67^. Small fragments of tissue-covered skeleton (< 2 cm^3^) were collected from each colony using a hammer and a chisel, and were preserved in 80% ethanol for further molecular analysis.

### Environmental conditions

The temperature and light intensity were monitored over a one-month period (i.e., from August 23 to September 26 2019) to assess environmental differences between the three reef habitats at Papetoai. Temperature was recorded for each habitat in 60-sec intervals using an *in situ* deployed HOBO Pendant Temperature Data Logger (Onset, USA), while the light conditions were recorded in 90-sec intervals using a 2π PAR Loggers (Odyssey, New Zealand). Differences in daily temperature and light intensity between reef habitats were assessed using one-way ANOVA or Kruskal-Wallis tests (in cases where ANOVA assumptions of normality and homoscedasticity were not met) with the package ‘stats’ implemented in the R software v3.6.3^68^, and the complemented post hoc pairwise comparisons were also conducted.

### DNA isolation and fire coral genotyping

The 198 colonies of *M.* cf. *platyphylla* were sampled and genotyped using microsatellite markers (as described in Dubé et al.^66^). Briefly, DNA isolation was performed using a QIAxtractor automated genomic DNA extraction instrument, according to manufacturer’s instructions, following an incubation at 55°C for 1 hr in 450 µl of lysis buffer with proteinase K (QIAGEN, Hilden, Germany). Each colony was genotyped using twelve microsatellite loci^69^. Samples were sent to the GenoScreen platform (Lille, France) for fragment analysis on an Applied Biosystems 3730 Sequencer with the GeneScan 500 LIZ size standard. All alleles were scored and checked manually using GENEMAPPER v4.0 (Applied Biosystems, Foster City CA, USA). Multilocus genotypes (MLGs) were identified in GENCLONE v2.0^70^. To assess the discriminative power of the microsatellite markers, the genotype probability (GP) was estimated at each locus and a combination of all loci in GENALEX v6.5^71^. Colonies with the same alleles at all loci were assigned to the same MLG (genet) and were considered as clone mates due to fragmentation. The 198 colonies belonged to ten different genotypes with at least four clonal replicates in at least two of the three surveyed habitats, provide an ideal study system to examine variation in Symbiodiniaceae communities among fire coral clones across distinct reef habitats (as a proxy for different environments) (Fig. 1, see Supplementary Data 1 for sample overview). A phylogenetic tree based on Nei’s genetic distances between the ten genotypes was inferred using the ‘poppr’ R package^72^ with bootstrap support from 5000 replicates (Supplementary Fig. 1A). A map with the x and y coordinates of each clonal genotype was produced using the R package ‘ggplot2’^73^ (Supplementary Fig. 1B).

### Symbiodiniaceae ITS2 library preparation, sequencing, and SymPortal analysis

The Symbiodiniaceae ITS2 region of the rDNA was amplified using the primers SYM_VAR_5.8S2 and SYM_VAR_REV^74^ with added sequencing adapters (forward: 5’– <u>TCGTCGGCAGCGTCAGATGTGTATAAGAGACAG</u>GAATTGCAGAACTCCGTGAA CC–3’; reverse: 5’–<u>GTCTCGTGGGCTCGGAGATGTGTATAAGAGACAG</u>CGGGTTCW CTTGTYTGACTTCATGC–3’; Illumina overhang adaptor sequences are underlined). Ten µl PCRs containing 1 µl of template DNA (10–50 ng of DNA from each fire coral sample) and 0.25 µM of each primer were run in triplicate per sample using the Multiplex PCR Kit (QIAGEN, Hilden, Germany). Thermal cycler conditions for ITS2 PCR amplification were 95°C for 15 min, followed by 30 cycles of 95°C for 30 s, 56°C for 90 s, and 72°C for 30 s, with a final extension time of 10 min at 72°C. Amplification success was verified by running 5 µl of the PCR products on a 1 % agarose gel, and successful triplicate reactions were pooled and cleaned using the Illustra ExoProStar PCR and Sequence Reaction Clean-Up Kit (GE Healthcare Life Sciences, Pittsburgh PA, USA). Indexing adaptors were added via PCR (8 cycles) according to the Nextera XT DNA library preparation protocol using the Multiplex PCR Kit. Indexed PCR products were purified and normalized using the SequalPrep Normalization Plate Kit (Invitrogen, Carlsbad CA, USA), subsequently quantified using the Qubit dsDNA HS Kit (Invitrogen, Carlsbad CA, USA), and run on the Agilent 2100 Bioanalyzer using the Agilent High Sensitivity DNA Kit (Agilent Technologies, Santa Clara CA, USA) to confirm amplicon length and purity. The ITS2 gene amplicon library was sequenced at the KAUST BioScience Core Laboratory on the Illumina HiSeq 2500 platform using the rapid-run mode with 2 x 250 bp overlapping paired-end reads with a 10% phiX control. Determined sequencing data for this project are available under NCBI BioProject <u>PRJNA888138</u>. Paired-end sequencing reads were submitted to SymPortal for quality control and analysis at SymPortal.org^75^. Briefly, sequence quality control was conducted as part of the SymPortal pipeline using mothur v1.39.5^76^, the BLAST+ suite of executables^77^, and minimum entropy decomposition (MED)^78^. ITS2 type profiles, representative of putative Symbiodiniaceae genotypes, were predicted by searching for co-occurring sets of sequences and characterized by specific sets of defining intragenomic ITS2 sequence variants (DIVs). Where ITS2 sequence abundances are reported in this study, we refer to post-MED, rather than pre-MED, abundances.

### Symbiodiniaceae community composition analysis

To determine the relative contribution of host genetic background (represented by the ten different host genotypes) and environment (represented by the three contrasting reef habitats) on coral-algae symbioses, two different methods of assessing the Symbiodiniaceae community composition were used: (1) ITS2 sequence assemblages were assessed as a proxy for total Symbiodiniaceae taxonomic composition and (2) predicted ITS2 profiles were assessed as a proxy for specific Symbiodiniaceae genotypes. As such, ITS2 sequence (post-MED) and ITS2 type profile abundance count tables (Supplementary Data 2 and 3), as output by SymPortal, were directly used to plot data using the relative abundances for each sample, as well as the presence or absence of a given sequence in a sample. Plots were drawn using ggplot2. To assess the explanatory power of host genotype and reef habitat variables on the variation in Symbiodiniaceae community composition, a two-way permutational multivariate analysis of variance (PERMANOVA) was run with the adonis2 function in the ‘vegan’ R package^79^. Dissimilarities between samples were visualized using a principal coordinate analysis (PCoA) using the ‘ape’ R package^80^. All statistical analyses were performed on Bray-Curtis distances of square root transformed ITS2 sequence counts using the vegdist function in vegan. We also analyzed our data to test for relationships between Symbiodiniaceae taxonomic composition dissimilarity (Bray-Curtis matrix) and host genetic distances (Euclidean matrix computed in GENODIVE v3.04)^81^ as well as host spatial distances (Euclidean matrix in meters computed in GENALEX v6.5) using Mantel tests and the Spearman correlation method with 9999 permutations using the function mantel in vegan. A hierarchical clustering analysis was performed on non-transformed ITS2 type profile counts using the function hclust in the R package stats to establish dissimilarity between the different identified *Symbiodinium* genotypes (i.e., sequences that belong to the *Symbiodinium* genus, formerly clade A^2^). Scripts used for analysis and plotting are deposited in a GitHub repository (https://github.com/CarolineDUBE).

## Data availability

Sequencing data determined in this study are available under NCBI BioProject ID <u>PRJNA888138</u>. Other data are available in the Supplementary Data files.

## Acknowledgements

We thank S. Planes for his contribution to the previous genotyping effort of *Millepora* samples at Moorea and KAUST Bioscience Core Lab for the ITS2 sequencing. This work was funded by the Laboratoire d’Excellence “CORAIL” project COMIC and KAUST baseline funds to CRV.

## Contributions

C.E.D and A.M collected the samples. C.E.D extracted the DNA and prepared the ITS2 libraries. C.E.D carried out the analyses with the contribution from B.C.C.H, A.M, and M.Z. C.E.D, E.B and C.A.-F.B conceptualized the study. C.E.D and C.R.V supervised the study. C.E.D, B.C.C.H and C.R.V wrote the manuscript with contribution from all the authors.

## Supplementary materials

**Supplementary Table 1.** Mantel tests showing the relationship between the dissimilarity of Symbiodiniaceae taxonomic composition (Bray-Curtis matrix) and spatial distance (Euclidean matrix in meters) for each host genotype (all clones combined irrespective of habitat) and genotype x habitat combination. Significant correlations are shown in bold.

| Genotype<br>(all clones across habitats) | Mantel<br>statistic (*r*) | p-values | min distance<br>(m) | max distance<br>(m) |
| --- | --- | --- | --- | --- |
| **G1** | **0.5658** | **0.001** | **1.05** | **188.60** |
| G2 | 0.1242 | 0.092 | 0.07 | 55.07 |
| **G3** | **0.3563** | **0.008** | **0.15** | **235.54** |
| **G4** | **0.7272** | **0.001** | **0.58** | **244.48** |
| G5 | 0.0924 | 0.238 | 0.90 | 228.28 |
| **G6** | **0.3538** | **0.002** | **0.07** | **192.04** |
| G7 | 0.0580 | 0.308 | 0.10 | 202.81 |
| **G8** | **0.3863** | **0.010** | **0.28** | **180.22** |
| G9 | 0.1069 | 0.237 | 0.22 | 231.04 |
| G10 | 0.0358 | 0.316 | 0.25 | 84.54 |
| Genotype x Habitat<br>(all clones in one single habitat) | Mantel<br>statistic (*r*) | p-values | min distance<br>(m) | max distance<br>(m) |
| G1 / BR | 0.4286 | 0.208 | 3.93 | 11.43 |
| G1 / UP | 0.0303 | 0.375 | 1.05 | 32.83 |
| G2 / UP | 0.1489 | 0.239 | 0.07 | 10.64 |
| G2 / MD | -0.2364 | 0.642 | 1.17 | 8.20 |
| G3 / BR | 0.0303 | 0.492 | 0.70 | 46.25 |
| **G3 / UP** | **0.2245** | **0.032** | **0.15** | **9.91** |
| G4 / UP | 0.6571 | 0.083 | 5.93 | 15.75 |
| G4 / MD | 0.0812 | 0.310 | 0.58 | 7.30 |
| G5 / UP | 0.0857 | 0.500 | 2.45 | 10.72 |
| G5 / MD | 0.0816 | 0.293 | 0.90 | 17.31 |
| G6 / BR | -0.1273 | 0.558 | 3.56 | 43.10 |
| **G6 / UP** | **0.5040** | **0.001** | **0.07** | **8.85** |
| G7 / BR | 0.8286 | 0.083 | 25.75 | 108.69 |
| G7 / UP | 0.1563 | 0.179 | 0.10 | 10.82 |
| G8 / BR | -0.1515 | 0.550 | 1.06 | 26.26 |
| **G8 / UP** | **0.6777** | **0.007** | **0.28** | **20.22** |
| G9 / UP | 0.0937 | 0.348 | 0.32 | 4.21 |
| G9 / MD | 0.0788 | 0.400 | 0.22 | 9.45 |
| G10 / UP | -0.0346 | 0.556 | 0.43 | 12.89 |
| G10 / MD | 0.3429 | 0.174 | 0.25 | 8.37 |

**Supplementary Figure 1.**
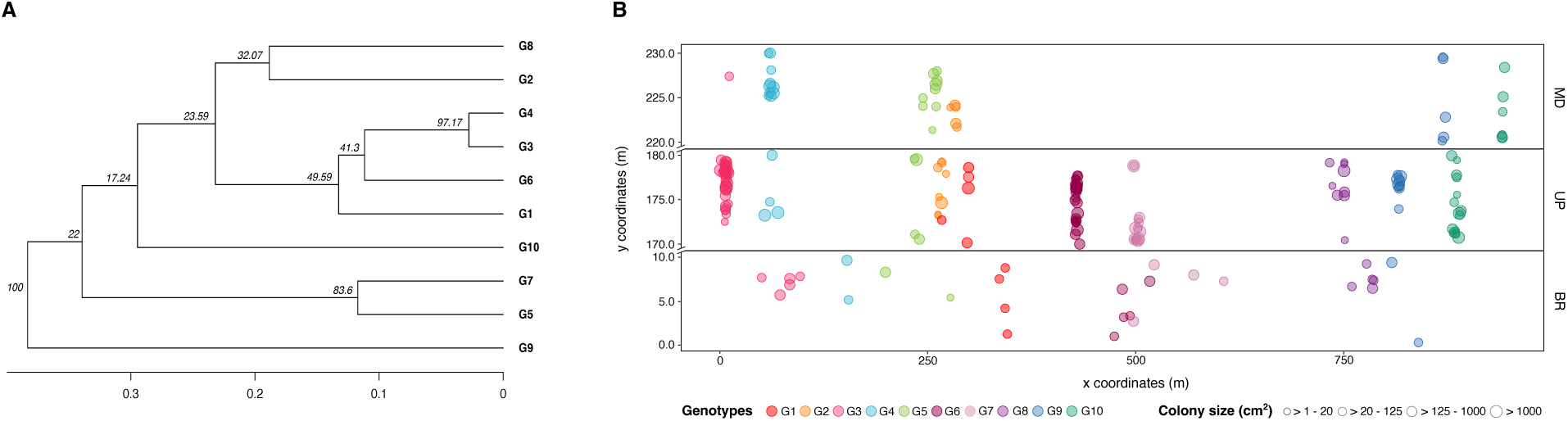
Genetic and spatial distances of fire coral genotypes. **A** Phylogenetic tree constructed by the neighbor-joining algorithm using Nei’s genetic distances. Numbers at the branch nodes indicate the confidence values for each branch obtained using the bootstrap procedure. **B** Map showing the spatial distribution of clones for each genotype. Points are plotted from belt-transects (300 m-long by 10 m-wide) and are shown on the x- and y-axis.

**Supplementary Figure 2.**
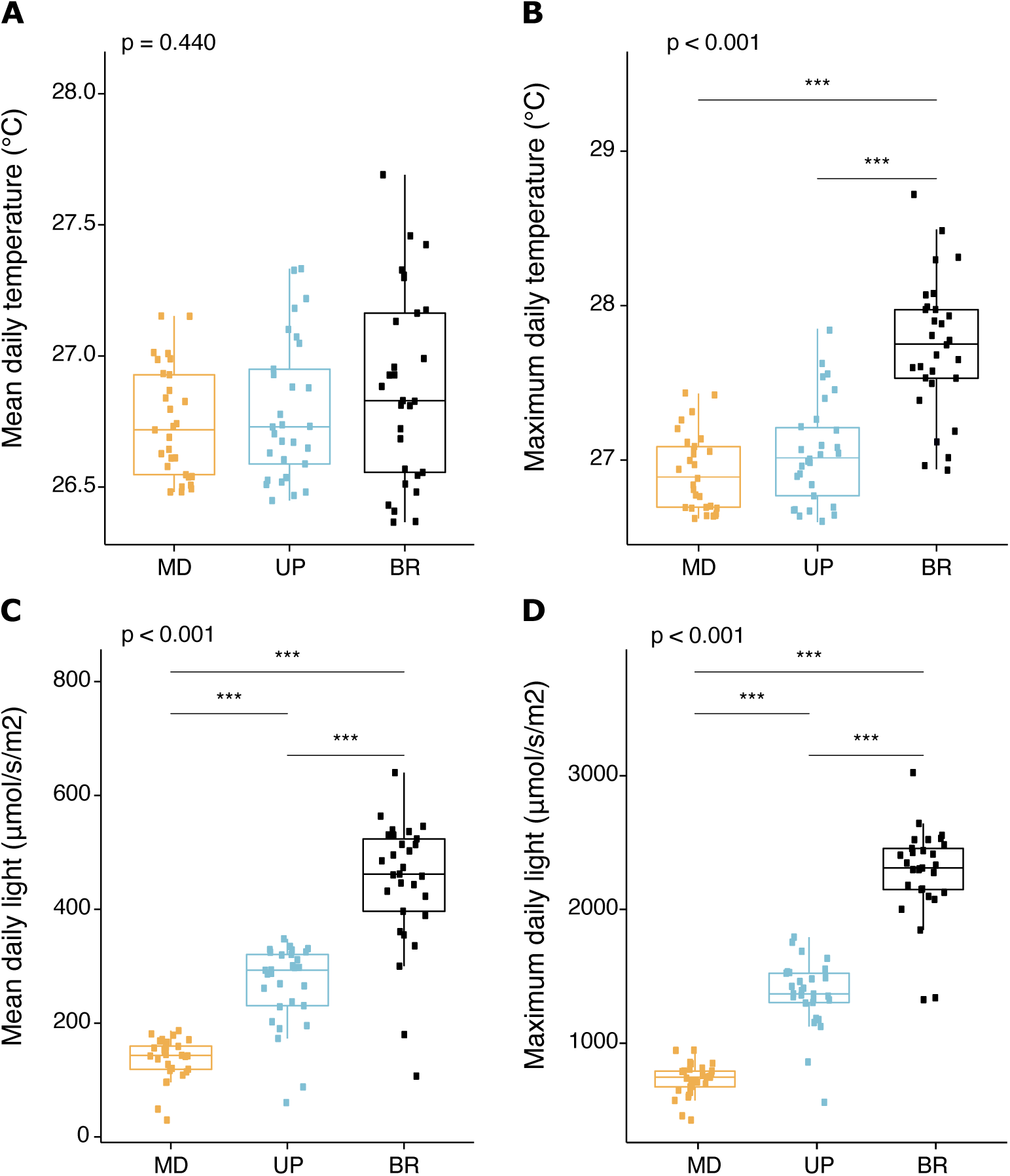
Environmental conditions of each reef habitat. **A** Mean temperature, (B) maximum temperature, (C) mean light, and (D) maximum light estimated at each of the habitat. The boxes represent the 25th to 75th percentile, lines show medians, and error bars depict 1.5X IQR. One-way Kruskal–Wallis test significance is shown on the top of each box plot (*P*-values), while post hoc pairwise comparison level *** refers to *P* < 0.001.

**Supplementary Figure 3.**
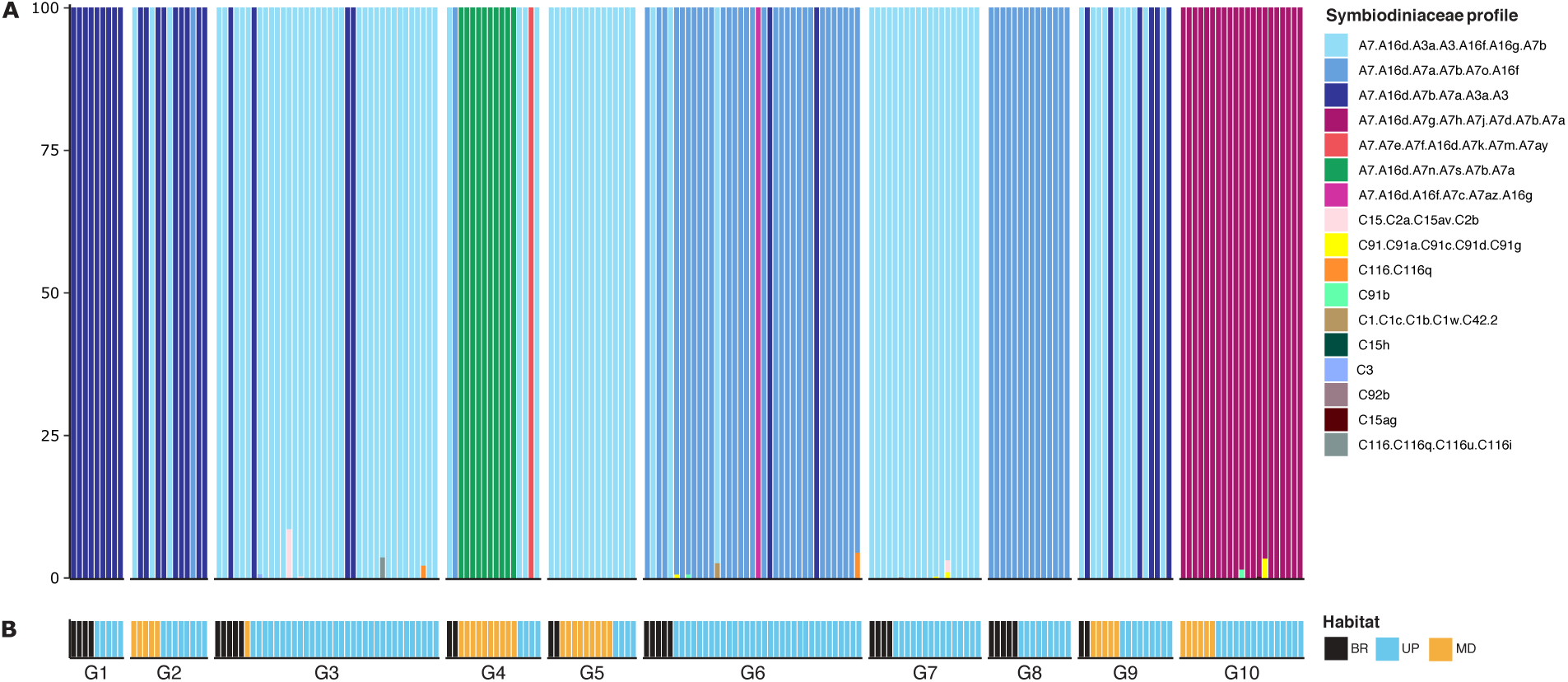
Symbiodiniaceae type profiles across clonal host genotypes. **A** Within each genotype, each sample is represented by a column of stacked bars showing the relative abundance of a given Symbiodiniaceae-type profile by identifying co-occurring sets of ITS2 intragenomic sequence variants. Profiles are annotated in the legend. **B** Each sample is represented by a column that defines its genotypic and environmental identity. Clonal genotypes are numbered from G1 to G10 and each genotype contains multiple samples (n) that occur naturally in at least two of the three surveyed habitats (BR = black, UP = blue, MD = gold).

**Supplementary Figure 4.**
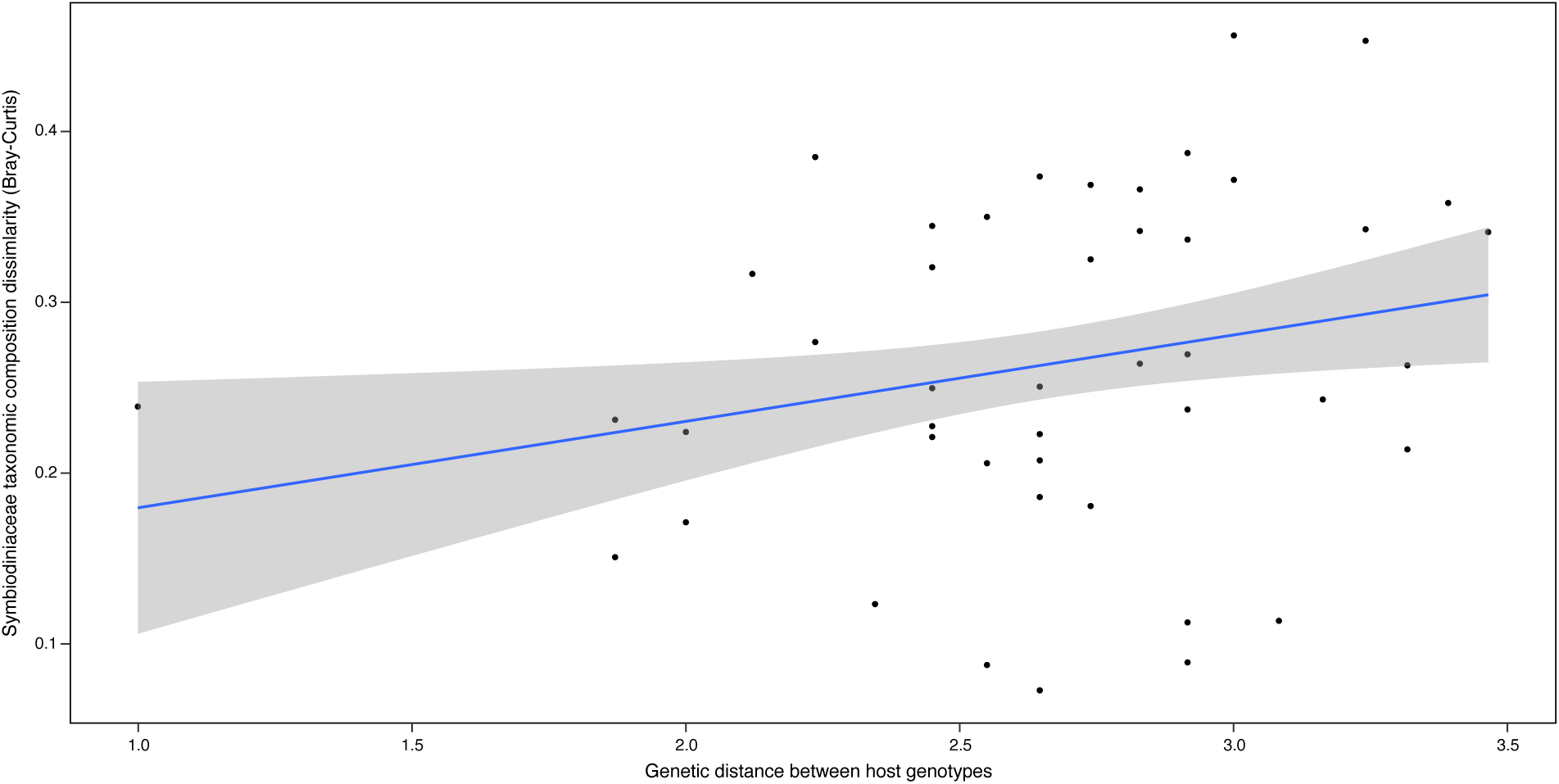
Relationship between *Symbiodinium* ITS2 sequence assemblage dissimilarity and genetic distance between host genotypes. Positive correlation between the relative abundances of *Symbiodinium* sequences based on Bray-Curtis distances and Euclidean genetic distances between fire coral genotypes. Clones of each genotype were grouped and the mean relative abundances of *Symbiodinium* sequences was calculated per genotype (Mantel test: *r* = 0.3258, *p* < 0.001).

